# Parallel evolution of *LRP2* gene mediating telescope-eye and celestial-eye in goldfish

**DOI:** 10.1101/2024.12.20.629592

**Authors:** Rongni Li, Bo Zhang, Yansheng Sun, Jingyi Li

**Author notes:** Corresponding author. Email: Jingyi Li.

## Abstract

After the intensive artificial selection, the development of celestial-eye in goldfish involves the protuberating and turning upwards of eyeballs, and also degeneration of the retinal. Thus, the celestial-eye goldfish provides an excellent model for both evolutionary and human ocular disease studies. Here, two mapping populations with segregating eye phenotypes in the offspring goldfish were constructed. Though whole genome sequencing using individual samples from the parents and pooled samples from the offspring, and RNA-seq for eyeball samples from pure goldfish lines, a premature stop codon in Exon 38 of *LRP2* gene was identified as the top candidate mutation that is responsible for celestial-eye in goldfish. Fatty acid metabolisms and epidermal cells, especially keratocytes related functions were inhibited in the eyeballs of celestial-eye, while inflammatory reactions and extracellular matrix secreting were stimulated. These suggest the dysfunction of cornea in the celestial-eye, and same for retinal, which could be the results of the truncated LRP2 protein. Besides, evidence was provided that not all the goldfish lines share the same causal mutation for celestial-eye, while the same gene *LRP2* is in charge of the similar phenotypes (celestial-eye and telescope-eye) in goldfish but no shared mutation. Therefore, those mutations and the associated phenotypes exhibit parallel evolutions in molecular level under artificial selections. Overall, the candidate mutation for celestial-eye in goldfish was identified by this study, and further analyses provide insights into the developmental and evolutionary processes of morphological changes in the eyes of goldfish.

**Author Summary:** As the first domesticated ornamental fish, goldfish is now worldwide spread and has undergone intensively artificial selections for morphological variations. Among them, celestial-eye is one of the most unique traits for goldfish, which consists of enlarged and protuberant eyes while the eyeballs turning upwards and thus suitable for top viewing in pots. In this study, we have identified that a single nucleotide mutation resulting in disrupted function of *LRP2* gene, is responsible for the celestial-eye phenotype. Together with our transcriptome analysis, the genetic and cellular mechanisms for celestial-eye in goldfish were reported for the first time, which provides fundamental knowledge for further studies about the development of eyes in fish and also for ophthalmic diseases in humans. In the aspect of evolution, our study and previous studies reported different truncated LRP2 proteins in charge of celestial-eye and telescope-eye, respectively, which are excellent materials to understand the mechanisms of parallel evolution in molecular level, i.e., independent artificial selections resulting in mutations of the same gene.

## Introduction

Goldfish is a variety of *crucian carp*. Beginning in the Song dynasty of ancient China (960-1,279 AD), probably from the lower Yangtze River [1], the goldfish have been intensively selected for their fascinating morphology traits [2]. Nowadays, numerous strains of goldfish with various shapes or colors serve as the world-famous ornamental fish and also excellent materials for understanding the genetics of morphology traits [3]. Among them, the strains with telescope-eye (TE) and celestial-eye (CE, also called as Sky Gazer, Heavenward Star Gazer, Chotengan, and Deme Ranchu) are popular strains among ornamental fish customers, while could also be utilized as the disease models for the eye development [4]. The TE phenotype consists of enlarged and protuberant eyes, while the CE goldfish possess the same character (**Fig 1**). However, the eyeballs of the TE goldfish are toward the side and front, like normal fish, while the eyeballs of the CE goldfish are upward, which are suitable for top viewing in pots. To be more specific, it is reported that the turning of the eyeballs beginning from 3 months of age to 4 months in the CE goldfish, maximum 6 months [5], while we observed that some of the CE goldfish completed their turning of eyeballs till 10 months of age, or even not able to turn their eyeballs to 90° upward at 24 months of age (**Fig 1**).

**Fig 1.**
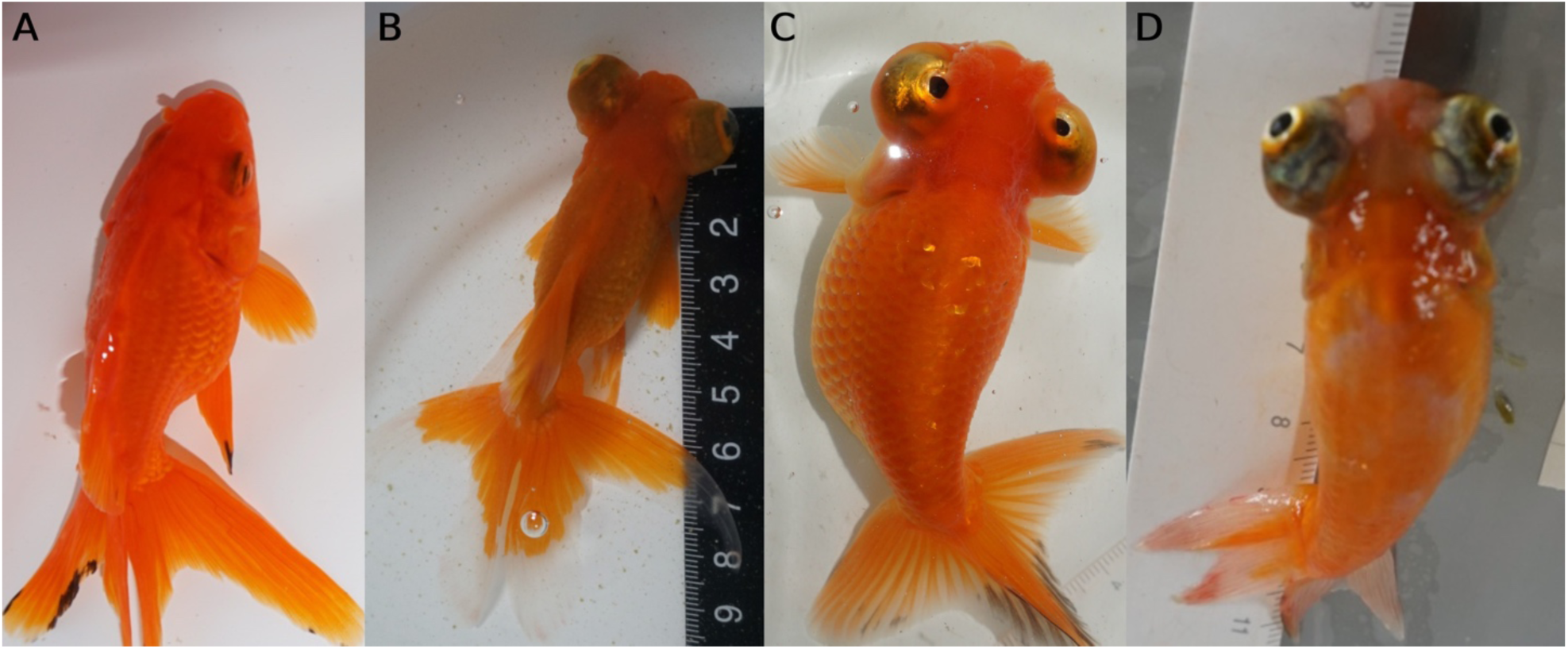
Variety of eye development in goldfish. (*A*) Goldfish with normal eyes (NE). (*B*) Goldfish with telescope-eyes (TE). (*C*) Goldfish with celestial-eye (CE). Turning angels of eyeballs were 90°. (*D*) Goldfish with CE. Turning angels of eyeballs were 45°. The above photos were taken by Rongni Li.

There has been long debate about the genetic relationship between the TE and CE goldfish. Considering their phenotypic similarity, it is reasonable to consider CE as a variety from TE with larger upward-turning eyes. However, according to the mitochondrial genome, Komiyama et al. have shown that TE and CE evolved independently, which took place in goldfish lineages with and without dorsal fin, respectively [6]. A possible earliest reference to the TE goldfish was recorded in 1590, while the CE goldfish appears to originated in the 18^th^ century [7]. Meanwhile, the possibility that gene flow allows the TE mutation to be exchanged into the goldfish without dorsal fin and contribute to CE could not be excluded. Yet, the allelic relationship between TE and CE was not reported yet. What is known is that they are both recessive to the normal eye (NE) goldfish [8].

The mutation of TE was denoted as *d* [8]. In 2020, Kon et al. reported three causal mutations of TE through GWAS, i.e., a retrotransposon 13 kb insertion in Intron 45 of *lrp2a* (*low-density lipoprotein receptor-related protein 2a*), and other two nonsense mutations for the same gene (Exon 51 and Exon 73). In addition, the gene editing of *lrp2a* via CRISPR/Cas9 which resulted in truncated proteins in two different goldfish strains (287 and 395 AA, respectively, compared with the original LRP2A protein with 4,653 AA), both created the TE phenotypes [10]. This provides compelling evidence that the incomplete form of LRP2A is responsible for TE in goldfish. However, it is unknown if TE and CE share the same causal mutation(s) at the molecular level.

Through the building of the mapping populations and a breed panel, diagnostic test, whole genome sequencing (WGS), and RNA-seq analysis, the current study revealed that the CE mutation affects the same gene but not sharing the same causal mutations with TE. Those findings provided clues to solve the previous debates, while also showing that under the artificial selections for CE and TE, parallel evolution occurred at the molecular level.

## Results

### Inheritance pattern of CE in goldfish

In a mating between CE goldfish, 1277 offspring were obtained including 1274 CE goldfish with various turning angles of their eyeballs, while only 3 were NE. The mapping populations were initiated by these 3 fish. Among them, one NE male was referred to NEM, the other two NE females were referred to NEF1 and NEF2.

The mating between NEM and NEF1 (Cross1) was carried out which produced 309 goldfish (**Fig 2**). Among them, 77 fish showed different degrees of CE (59 fish were 90° turned upwards, 12 fish were 80-90°, 4 fish were 45-80°, and 2 fish were 15°-45°), and the other 232 fish were NE. The numbers of CE and NE fish in the offspring of Cross1 matches 1:3 ratio (*P* > 0.05). In the mating between NEM and NEF2 (Cross2), 634 goldfish were obtained including 109 with CE (**Fig 2**). All these CE fish showed large turning angles of their eyeballs (64 fish were 90° turned upwards, 32 fish were 80-90°, 13 fish were 45-80°). Other than CE, 126 fish with slightly TE phenotype were also observed in Cross2. However, they failed to develop compete TE as their eyes were slightly protuberant without turning upwards. Those unexpected TE fish were excluded from our analysis. The other 399 fish in the offspring of Cross2 were NE. Thus, the CE and NE fish in Cross2 also match 1:3 ratio (*P* > 0.05). This data suggests that a single autosomal recessive gene is responsible for CE in our mapping populations.

**Fig 2.**
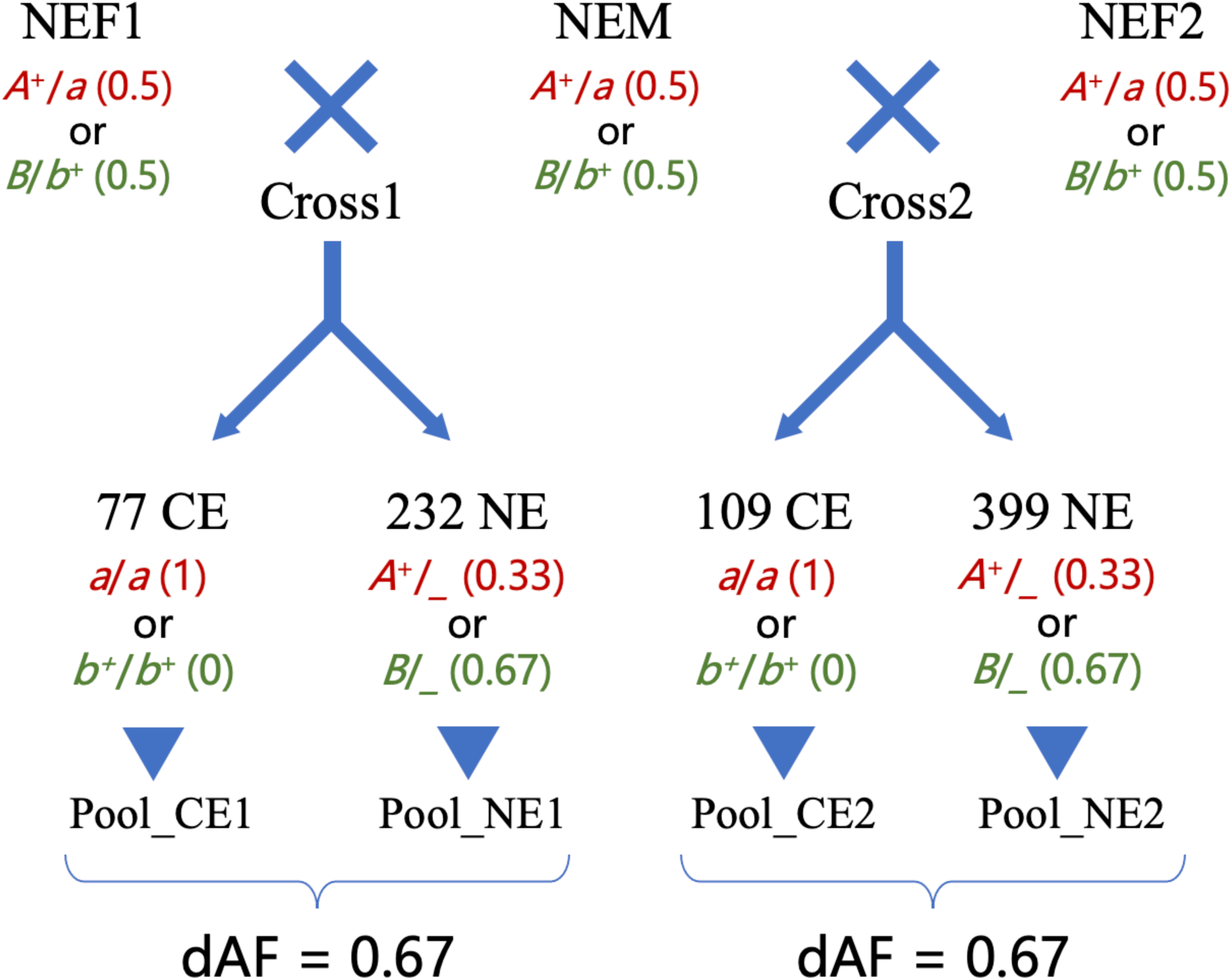
Schematic diagram of the mapping populations. NEF1, NEF2, NEM are the 3 parental goldfish with NE, while F and M refers to female and male, respectively. The two matings produced 77 offspring with CE, 232 with NE, 109 with CE, and 399 with NE. The four DNA pools were constructed from them. The red letters describe the assumption that a recessive mutation (*a*) causing CE segregated in the offspring, while the green letters are for the different possible scenario that a dominant inhibiting mutation (*B*) for CE segregated in the offspring. The numbers in brackets indicate the allelic frequencies of the mutant allele (*a* or *B*). dAF stands for differences of allelic frequencies between two pools.

### Chromosome 9 is associated with CE

For samples from the offspring of Cross1, 2 DNA pools were constructed based on the shared phenotypes (Pool_CE1, pooling of all 59 CE fish with eyeballs 90° turned upwards, and Pool_NE1, pooling of 80 randomly selected NE fish). Another 2 DNA pools (Pool_CE2, n = 64 and Pool_NE2, n = 80) were also constructed using samples from Cross2 with the same criteria of selecting samples for pooling. A total of 1.9 billion reads were obtained by sequencing these 4 pools and the average mapping rate is 99.34%. The 3 parental goldfish with NE were also sequenced individually, which generated 532 million reads with 99.30% of average mapping rate.

Since CE is recessive and the reference genome that we used came from a goldfish with NE [11], such putative mutation was denoted by *a* allele in our mapping populations. Thus, the mutant allele (*a*) should be fixed in Pool_CE1 and Pool_CE2 (as red fronts in **Fig 2**).

Alternatively, the entire mapping populations could be fixed for the CE mutation but a dominant inhibitor mutation (denoted by *B* allele in **Fig 2**) for CE was segregated. In this case, the wild-type allele (*b^+^*) should be fixed in Pool_CE1 and Pool_CE2 (as blue fronts in **Fig 2**). Under these two assumptions, the theoretic allelic frequencies of the mutant allele for each of the parent should be 0.5; for Pool_CE should be 1 or 0, respectively; for Pool_NE should be 0.33 or 0.67, respectively (since in the offspring, 25% were homozygous mutants, 50% were heterozygotes, and 25% were homozygous wild-type).

By comparing Pool_CE1 and Pool_NE1, regions with ZF_ST_ larger than 11 were found in chromosome 9 (7 regions with the total size of 2.92 Mb, from 21.39 Mb to 28.63 Mb, see **Fig 3 and S1 Table**, the coordinates refer to the goldfish genome assembly reported by Chen et al., in 2020, the same below). A total of 8,361 SNPs and 2,876 other types of variants were defined as candidate mutations by only considering the ones with allelic frequency differences (dAF) between the 2 pools larger than 0.5 (As mentioned above, the dAF should be 0.67 in theory (**Fig 2**). In case of pooling or sequencing error, the threshold of dAF was set as 0.5) while were heterogeneous in NEM, NEF1, and Pool_NE1, and also were homozygous mutant or wild-type in Pool_CE1.

**Fig 3.**
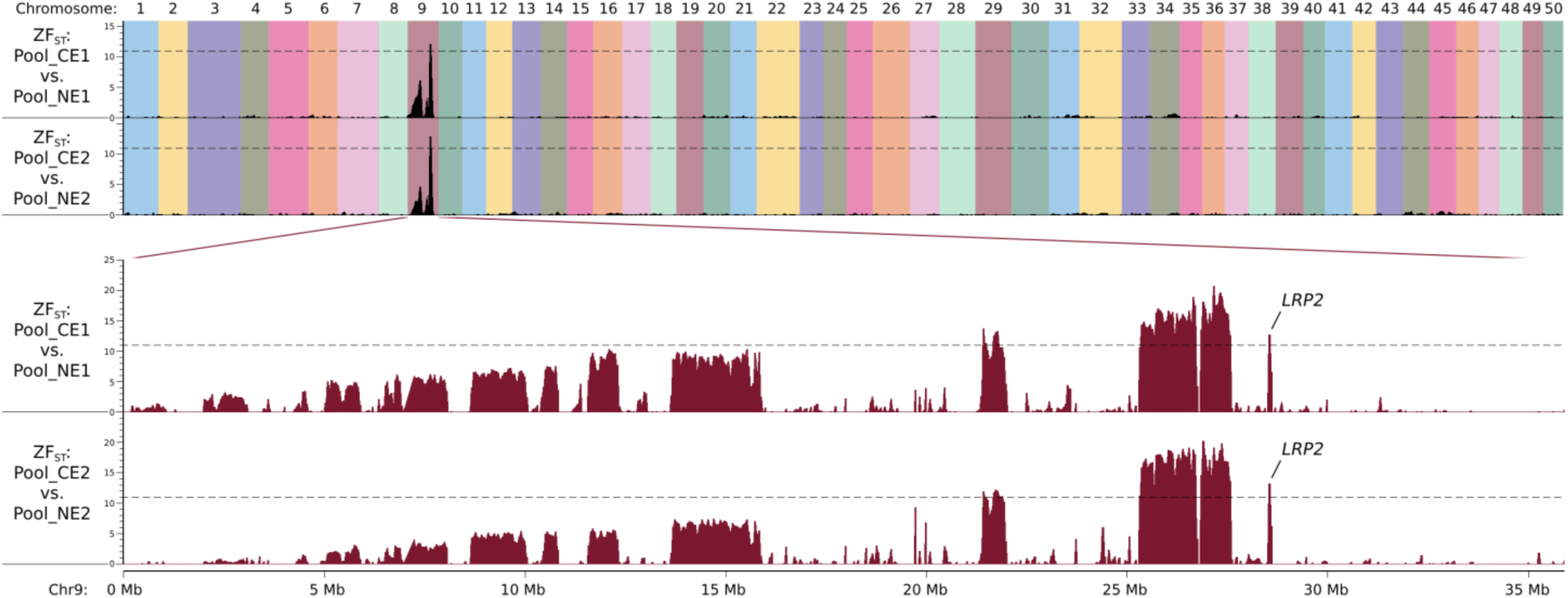
Comparisons of DNA pools revealed selective regions in Chromosome 9 associated with celestial-eye (CE) in goldfish. Pool_CE1 and Pool_NE1 consist of 59 CE and 80 NE (normal eye) fish in the offspring of the first family, respectively. Pool_CE2 and Pool_NE2 consist of 64 CE and 80 NE fish in the offspring of the second family, respectively. Regions with ZF_ST_ score larger than 11 (dash lines) were considered as candidate regions. Chromosome 9 was roomed-in in the lower panels. The location of *LRP2* gene was indicated.

By comparing Pool_CE2 and Pool_NE2, with the same criteria, the candidate region was defined as 8 regions on chromosome 9 (with the total size of 3.05 Mb, from 19.73 Mb to 28.63 Mb, see **Fig 3 and S1 Table**, discontinuous candidate regions for both Cross1 and Cross2 could be due to miss assembly of the reference genome) and 8,368 SNPs and 2,874 other mutations were selected for further analysis. The two sets of candidate regions defined by these two comparisons largely overlap (**Fig 3**), suggesting that NEF1 and NEF2 share the same causal mutation, and should be also the same with NEM.

### Known TE causal mutations were not detected in the CE goldfish

Since the regions associated with CE (19.73∼28.63 Mb) harbors previously reported TE causal mutations, we firstly investigated those mutations in our populations. Those mutations include the two nonsense mutations in the exons 51 and 73 of *lpr2aL* [9,10] (corresponding to chr9:28,593,475 and 28,616,634 in the goldfish genome assembly reported by Chen et al., in 2020), and the ∼13 kb retrotransposon insertion in the intron 45 of the same gene [9]. As the results, the two nonsense mutations were not detected in the mapping populations in this study, while there are five SVs in the intron 45 of *lpr2aL* by viewing the BAM files. Two of them are microsatellite variations, another two could be transposon elements (**S1 Fig**). Thus, primers were designed and Sanger sequencing were applied to further confirm the existence of the ∼13 kb insertion in our CE goldfish. In the 4 CE offspring from our mapping populations, no retrotransposon specific fragment was amplified. However, the amplification of the entire intron 45 produced a ∼2.5 kb fragment which is although ∼0.8 kb longer than the reference sequence but cannot harbor the complete ∼13 kb insertion. The sequencing of such PCR products revealed that the leftmost SV in **S1 Fig** is actually a 33 bp sequence (chr9:28,583,267-28,583,299) replaced by a 116 bp sequence. Other SVs were unable to be Sanger sequenced because of the microsatellites. However, it is likely that those SVs also added hundreds of base pairs which contributed to the ∼0.8 kb extra sequence. In conclusion, the ∼13 kb insertion is not detected in the CE goldfish. Altogether, none of the known TE causal mutations was detected in our CE mapping populations.

### CE in goldfish is heterogeneous

Outside of our mapping populations, 50 CE goldfish from 5 different fish farms were whole genome sequenced individually, which generated 3.2 billions of reads with 99.38% of average mapping rate.

In the assumption that a single mutation caused all the CE phenotypes, we firstly focused on the mutations that were fixed in all the 52 CE goldfish libraries (Pool_CE1, Pool_CE2, and the 50 CE individuals) from our candidate mutations. As a result, 19 mutations were screened out. However, the majority of them have low calling rate since quality control for the mutations was not applied. Therefore, after manually confirming their genotypes by viewing the BAM files, the majority of the allelic frequencies were corrected. As the results, only 2 SNPs matched the criteria (fixed in the 52 CE goldfish libraries while heterogeneous in other 5 libraries) and underwent further investigation. These 2 SNPs (chr9:26,872,019 and 26,872,024) are 5 bps far away, located 100 kb and 39 kb downstream of *retsatl* and *Cau.09G0011130*, respectively. They were homozygous WT in all the CE goldfish libraries so that the mutant allele may act as an inhibitor of CE, if they are functional. They were also possible no function variants but merely carried by the 3 NE parents, since they are downstream mutations and far away from the genes.

Therefore, we considered the alternative assumption that not all the 50 CE goldfish outside of our mapping populations carried the same CE causal mutation. In order to determine which samples and which regions may harbor the same causal mutation with our mapping populations, pair-wise genetic distances between Pool_CE1 or Pool_CE2 and each of the 50 individual samples were calculated for each candidate region (**Fig 4**). Within each region, the individual samples with any genetic distance larger than 0.1 were excluded for the target mutation screening (**S2 Table**), and the criteria is similar that target mutations should be fixed in all the remaining CE libraries. As the results, 6 out of 9 candidate regions (Candidate regions 1 to 6 in **S1 Table**) were excluded. Although there are still a large number of target mutations (10,146), they are located within a ∼3.2 Mb region (chr9:25,346,752-28,589,750, defined as the target region, including 59 annotated genes). It is also clear that not all the CE goldfish shared the same IBD (Identity-by-descent) sequence in the target regions (**Fig 4**).

**Fig 4.**
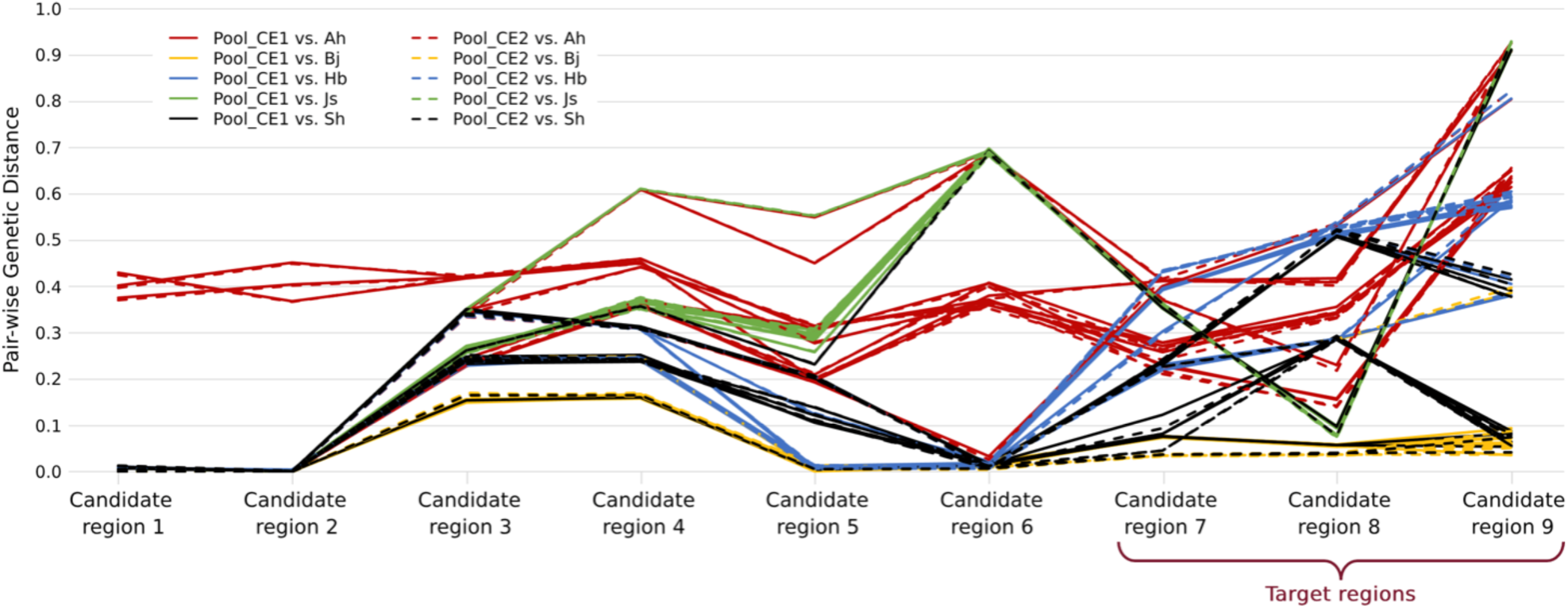
Pair-wise genetic distances between the DNA pools from the mapping populations and individual samples outside of the mapping populations. The DNA pools are Pool_CE1 and Pool_CE2, which are the offspring expressing celestial-eye (CE) in the mapping populations. The individual CE samples are from 5 different fish farms located in different regions in China, i.e., Anhui (Ah), Beijing (Bj), Hebei (Hb), Jiangsu (Js), and Shanghai (Sh) (n=10 for each farm). Candidate regions were previously defined by ZF_ST_ values. After excluding the samples with genetic distances larger than 0.1 for each region, the target mutations (fixed in all the remaining samples) only distributed in candidate regions 7 to 9, and thus were termed as target regions.

### Putative functional mutations were identified as candidates for CE

In case of artifacts during sequencing, all the SVs in the expanded region, from 24.35 to 29.59 Mb in chromosome 9 (1 Mb upstream and 1 Mb downstream of the target regions) were investigated for their putative functions. Under the combination of Lumpy software and manually double-checking, 97 SVs were detected from the 7 libraries of the mapping population (3 parental samples and 4 offspring pools). Eventually, 2 SVs (a 1.5 kb deletion and a 200 kb complex SV possibly involved deletion and inversion) were sifted out. Next, the genotypes of these 2 SVs in the 50 individual CE samples were determined by viewing the BAM files, which showed that these 2 SVs still match the criteria of candidates according to **S2 Table** (in Candidate regions 7 and 8, respectively). This 1.5 kb deletion (chr9:25,574,191-25,575,687) is 3.8 kb downstream of an uncharacterized coding gene (Cau.09G0010620) and 36.2 kb downstream of *CALCRLA*; the 200 kb SV (and 27,182,149-27,382,584) harbored 3 annotated genes (*HDAC4*, *TRAF3IP1*, *TWIST2*).

Among the target mutations (SNPs and small Indels, n=10,146, as defined above), there are 11 frameshift Indels, 6 non-frameshift but coding Indels, 119 nonsynonymous, 1 stopgain, and 1 stoploss SNPs. Those mutations involve 18 other annotated genes. Together with the 4 genes probably being affected by the 2 candidate SVs, these 22 genes are defined as candidate genes.

### Epidermis related processes, fatty acid metabolisms, and immune responses were involved in the formation of CE

In order to further narrow down the candidate genes in the target regions, RNA-seq was applied using eyeball samples from NE and CE goldfish (14 months of age). As the heatmap shows, the RNA-seq samples cluster according to their phenotypes, NE or CE (**Fig 5A**). A total of 4,665 differentially expressed genes (DEGs) were detected. Among them, 2,499 were down regulated and 2,166 were up regulated in the CE group (**Fig 5B**).

**Fig 5.**
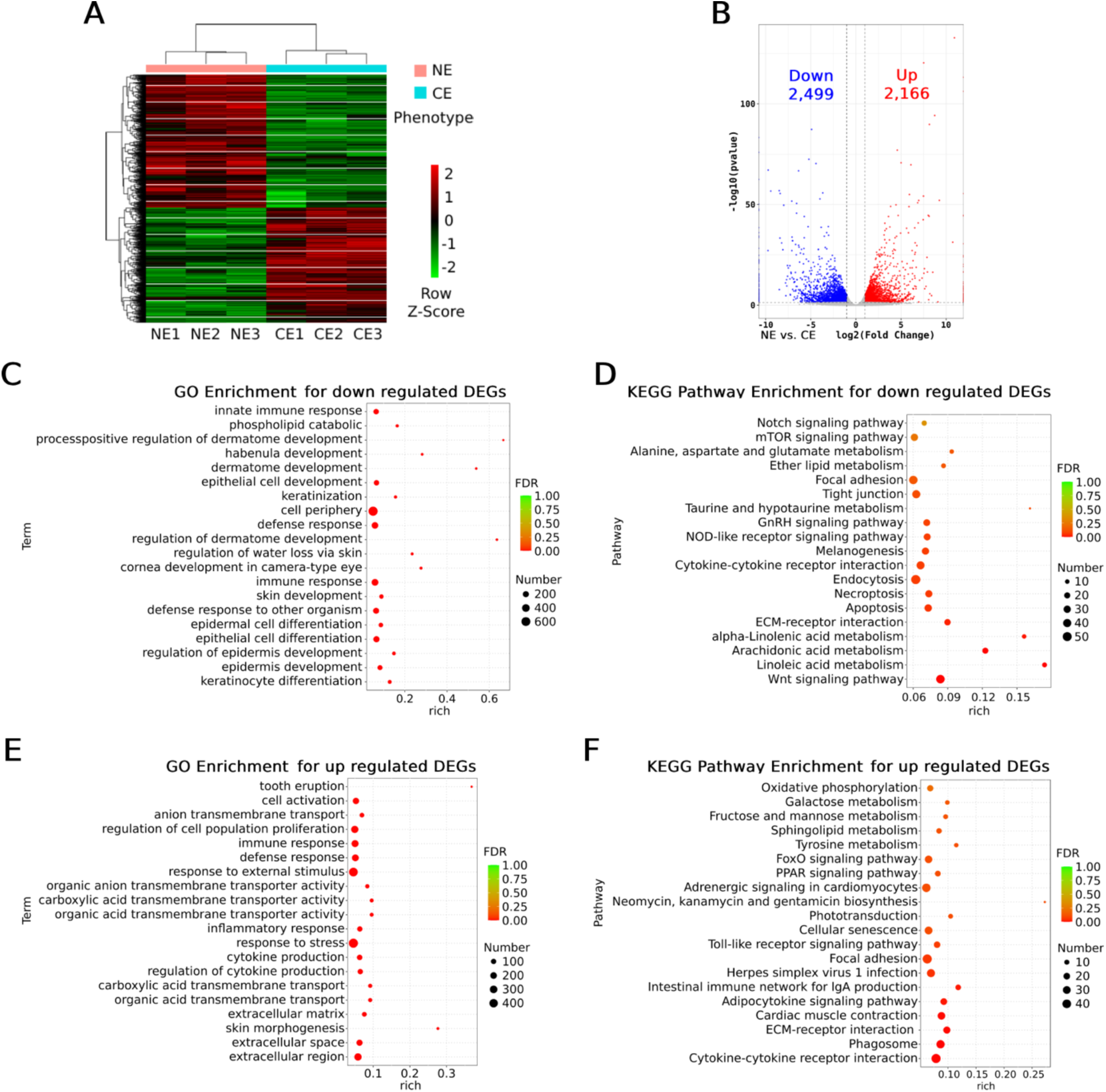
Comparison of expression profiles between eyeballs of NE and CE goldfish via RNA-seq. NE stands for normal eye and CE stands for celestial-eye. (*A*) Heatmap for the gene expressions in the 3 NE samples and 3 CE samples. (*B*) Volcano plots showing the differentially expressed genes (DEGs), including up and down regulated genes in CE samples. The criteria for DEGs are |log2foldchange| > 1 and adjusted *P*-value < 0.05. Top 20 GO terms (*C*) and top 19 KEGG pathways (*D*) significantly enriched (*P* < 0.05) for the DEGs down regulated in CE samples. Top 20 GO terms (*E*) and top 20 KEGG pathways (*F*) significantly enriched (*P* < 0.05) for the DEGs up regulated in CE samples. In each of *C* to *F*, the *P*-value increases as the terms or pathways are presented from the bottom to the top.

The down and up regulated DEGs were respectively applied to the enrichment analysis by Gene Ontology (GO) and Kyoto Encyclopedia of Genes and Genomes (KEGG) pathways. For down regulated DEGs, all the top 5 most significant enriched GO terms are epidermal cell related processes (keratinocyte differentiation, epidermal cell development, and differentiation, **Fig 5C**), while 3 out of the top 4 most enriched KEGG pathways are fatty acid metabolisms (linoleic acid and arachidonic acid metabolisms, **Fig 5D**). For up regulated DEGs, 6 out of the top 10 most enriched GO terms are also epidermal cell related or extracellular terms (also considered as epidermis related, including extracellular region, space, and matrix, and response to external biotic stimulus, **Fig 5E**), in the rest of 4 GO terms, 3 are immune response related (immune response, defense response, immune system process, ranking 2^nd^ to 4^th^ most enriched terms, **Fig 5E**). Immune response pathways were also enriched as the top pathways in the KEGG analysis for up regulated DEGs (6 pathways out of the top 10, cytokine-cytokine receptor interaction, phagosome, adipocytokine signaling pathway, intestinal immune network for IgA production, herpes simplex virus 1 infection, and Toll−like receptor signaling pathway, **Fig 5F**), suggesting that inflammatory reactions may be taken place in the eyeballs of the CE goldfish. Besides, PPAR (peroxisome proliferator-activated receptor) signaling pathway was significantly enriched for the up regulated DEGs (**Fig 5F**), which is also key for fatty acid metabolism. In addition, terms of cornea development in camera-type eye and regulation of water loss via skin were enriched for down regulated DEGs, together with keratinocyte related terms (**Fig 5C**), suggesting dysfunction of cornea in the CE goldfish; melanogenesis pathway was significantly down regulated (**Fig 5D**), while pathways of tyrosine metabolism and phototransduction were up regulated in the CE goldfish (**Fig 5F**), suggesting that retina was also affected. When applying enrichment analysis for all the DEGs, terms including skin morphogenesis, inflammatory response, and other extracellular related terms, pathways involving immune response and fatty acid metabolism, were also enriched (**S2 Fig**).

Taken together, our RNA-seq data reveals that epidermis related functions including extracellular processes were dramatically changed in the eyeballs of the CE goldfish, while fatty acid metabolisms were inhibited, and immune responses especially inflammatory reactions were stimulated. Functionally important genes in cornea and retina could be differentially expressed.

### *LRP2* and its coding mutations are the top candidates

Among the 59 annotated genes in the target regions, 9 of them were differentially expressed according to our RNA-seq data. They are *CERKL*, *NEUROD1*, *ITPRID2*, *FRZB*, *AGR3*, *LOC113071285*, *KRT18*, *klhl41a*, and *LRP2* (**Fig 6A**). To understand which DEGs could be more upstream regulators to others, their interaction network and gene prioritization were predicted (**Fig 6B**). As a result, *NEUROD1*, *FRZB*, *KRT18*, and *LRP2* could play more central roles rather than *ITPRID2* and *AGR3*. *CERKL* is predicted to be independent from the network of **Fig 6B**, while *LOC113071285* and *klhl41a* were not included in the database of GeneMANIA.

**Fig 6.**
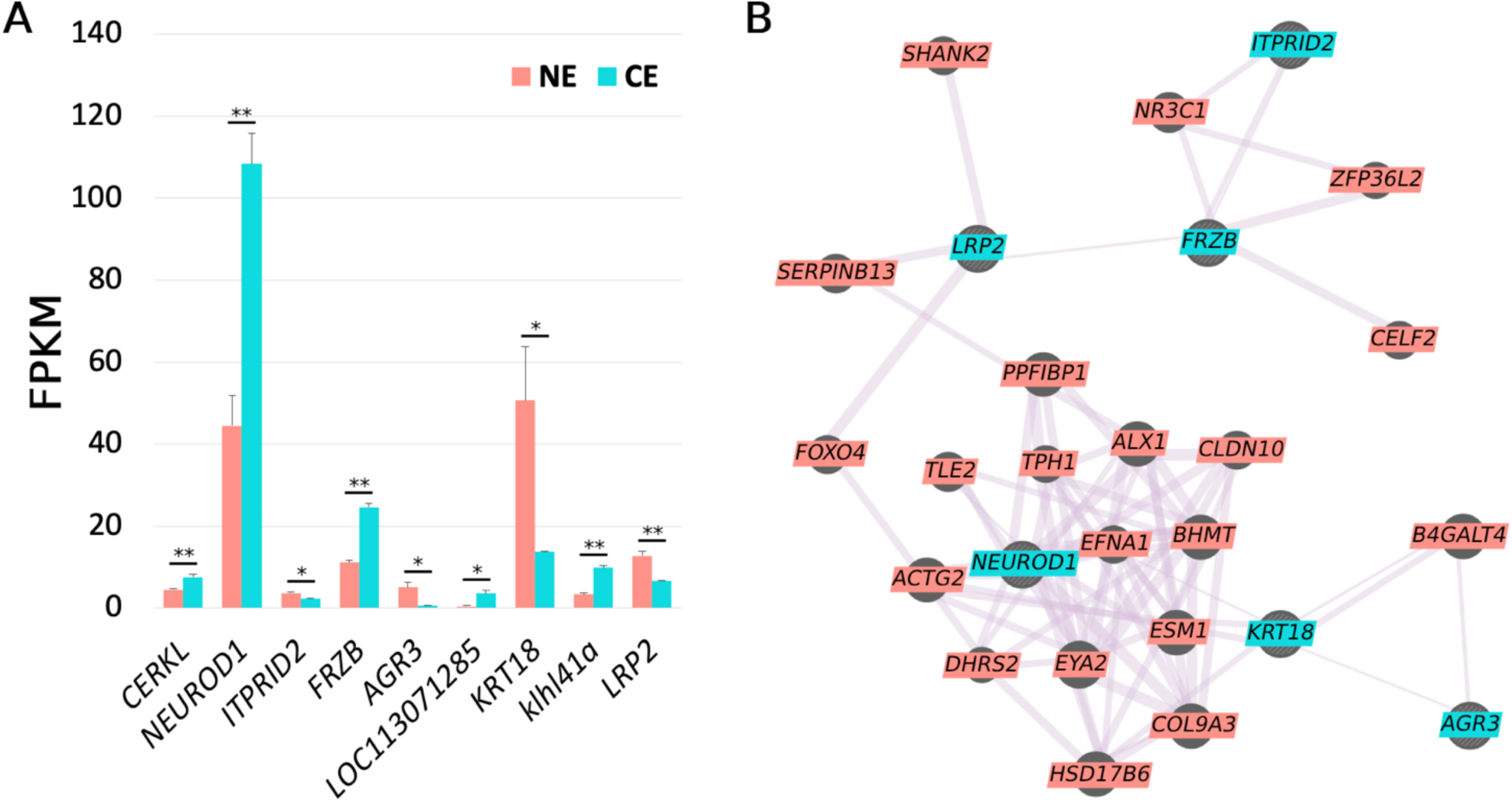
Differentially expressed genes (DEGs) located in the candidate regions of CE in goldfish. (*A*) Within the candidate regions, 9 DEGs were detected via RNA-seq. NE stands for normal eye and CE stands for celestial-eye. Data are represented as mean ± SEM. An asterisk indicates significant difference (P < 0.05) while double asterisks indicate extremely significant differences (P < 0.01). Total RNA was extracted from eyeballs, n = 3 for each column. (*B*) Predicted gene interactions for the DEGs (blue color, 3 DEGs were not listed since no data or no interaction) and other related genes (pink color) according to the GeneMANIA database.

Among the 22 candidate genes putatively affected by coding mutations or candidate SVs, 4 of them are DEGs: *CERKL*, *ITPRID2*, *FRZB*, and *LRP2*. None of them was adjacent to the SVs. Since the 3 NE goldfish used for RNA-seq came from a different population than our mapping populations, we checked the BAM files for the 3 RNA-seq data to genotype those putative functional mutations. We found that, for the 2 nonsynonymous mutations of *CERKL* (chr9: 26,383,105 and 26,383,771), all the NE samples were heterozygotes. No reads were found for the other 2 nonsynonymous mutations of *CERKL* (chr9: 26,413,943 and 26,426,046) in NE samples. For *ITPRID2*, heterozygotes were found in 1 of the NE samples for the 2 frameshift mutations (chr9: 26,464,773 and 26,464,774) and in all the NE samples for the nonsynonymous mutations (chr9: 26,459,566). Only 2 reads were aligned to 1 of the NE samples showing the other frameshift mutation of *ITPRID2* (chr9: 26,464,066), and they were mutant alleles, other 2 NE samples were unknown for this mutation since no reads were found. For the nonsynonymous mutation of *FRZB* (chr9: 26,673,148), 1 NE sample was heterozygote, others were homozygous mutants. For the nonsynonymous mutation (chr9: 28,575,713) and the stopgain mutation (chr9: 28,575,379) of *LRP2*, the genotypes of all the 6 RNA-seq samples (the 3 CE samples were also checked) matched the candidate pattern, which is homozygous mutant in CE samples and homozygous wild-type in NE sample. Considering that those no call mutations could be tightly linked with other mutations in the same gene, or not functional since not expressed, *CERKL*, *ITPRID2*, and *FRZB* were excluded from candidate genes since heterozygous mutations were detected in NE samples. Those samples came from a population which is breed true for NE, so there should not be carriers for the CE causal mutation. At least, no other obvious functional mutations were found affecting those 3 genes. Thus, *LRP2* remained to be the only candidate gene. Additionally, we also analyzed the RNA-seq data reported by Du et al [12]. Comparing the whole embryo between CE and NE goldfish, LRP2 was found to be differential expressed in the 14-somite stage (expression in NE is 1.7-fold higher than that in CE, *P* < 0.01) but not zygote or 35%-OVC stages.

In summary, by analyzing the expression profiles and coding mutations of the RNA-seq data, there is multiple evidence supporting that *LRP2* and its 2 coding mutations (both in Exon 38) are the top candidates for CE in goldfish. Obviously, the premature stop codon of *LRP2* (chr9: 28,575,379) is more likely to be functionally important.

### The CE goldfish with different turning angles of eyeballs are associated with chr9:28,575,379

In our mapping populations, only a proportion of the offspring were sequenced as they were either CE with the eyeballs were 90° turned upwards, or NE. To investigate whether the premature stop codon in Exon 38 of *LRP2* is also responsible for the CE goldfish with different angles of eyeballs turning, 22 of the offspring in our mapping populations with various phenotypes were selected for genotyping the SNP of chr9:28,575,379 (C>T). As the results, in the goldfish with obvious protuberant eyes, no matter what angles of eyeballs turning (from 15° to 90°), or whether asymmetric of two eyeballs (e.g., a CE goldfish with one eyeball 90° turned upwards and one 60° turned), all of them were homozygotes of mutant alleles. Other goldfish with slightly protuberant eyes (no turning of eyeballs), or NE, were all homozygotes of wild-type allele or heterozygotes. Therefore, we proposed that, the premature stop codon in Exon 38 of *LRP2* (chr9:28,575,379) could be responsible for the protuberant and the upward turning of eyeballs in goldfish, while the angles of eyeballs turning were affected by other factors. The fact that the CE goldfish with various turning angles of eyeballs were homozygous of the CE mutation, is consistent with our deduction in our mapping population: the CE homozygous offspring were one fourth of all the offspring, in both two mapping populations, suggesting that a single autosomal recessive gene is responsible for CE and the three parental fish were heterozygous for this mutation.

## Discussion

In this study, an SNP was identified as the putative causal mutation for CE in goldfish, through the construction of phenotypic segregating populations, detection of selective regions through the whole genome, comparative genomics between different populations, transcriptomic analysis, and diagnostic test. This SNP will lead to a truncated protein of LRP2, which clarified the genetic relationship between the TE and CE goldfish. The truncated LRP2 were also reported to be the molecular mechanisms of TE in nature [9,10]. This could explain the phenotypic similarity between TE and CE. Meanwhile, no mutation was shared between TE and CE, showing that they evolved independently. This is consistent with the analysis of mitochondrial genomes of the TE and CE goldfish [6]. Therefore, we discovered the parallel evolution in molecular level between the TE and CE goldfish since different mutations in the same gene cause similar phenotypes. Moreover, we have shown that not all the CE goldfish were homozygous mutant at chr9:28,575,379 (in Candidate region 9 of **Fig 4**, indicating that most Bj and Sh individuals were homozygous mutant), there must be other causal mutation(s) for CE and thus other parallel evolutionary event(s) that were awaiting to be identified.

To be more specific, the LRP2 proteins without the 3’ portions coded by exons after intron 45, exon 51, or exon 73 lead to the TE phenotype [9,10], while the truncated LRP2 protein missing the 3’ portions coded by exons after exon 38 is associated with the CE phenotype. This suggests that the translation of exons 38 to 45 could be necessary for preventing the turning of eyeballs while the more downstream exons are responsible for whether the eyeballs were protuberated. However, these LRP2 proteins were fund in nature. The artificially edited goldfish that only exons 1 to 8 were translated into the LRP2 protein, showed the protuberated but not turned eyes [10]. The relationships between those exons and the phenotypes needs further investigation. These different truncated proteins provide excellent materials for understanding the detailed molecular mechanisms of LRP2.

In the cell level, the clinic features of the celestial eyeballs of goldfish have been carefully studied [4,5]. Here we can compare them with the molecular features acquired from the current study. At the same age (∼90 days), the eyeballs of the CE goldfish started to protrude and the retina started to degenerate, while the retina in the TE goldfish with protruding eyes did not obviously degenerate [5], although the TE goldfish has thinner retina [13]. This suggests that there is no causal relationship between eye expansion and retina degeneration, instead, they are both direct consequences of the CE mutation at the same time. Therefore, the two developmental processes are discussed separately:

Regarding the eye expansion, it might begin with modified lipid storage and metabolism in the eyeballs of the TE goldfish by over-expression of genes in PPAR signaling pathway which could transport fatty acids outside of the eyeballs [13]. It could be similar in the CE goldfish since the upregulation of PPAR signaling pathway and down regulation of fatty acid metabolisms were observed as well in the current study. Next, the altered lipid and fatty acid content in the eyeballs could be the cause of vitreous expansion and thus elevated intraocular pressure in the TE goldfish [14], and possibly in the CE goldfish. Regarding to the retina degeneration, at later age (120 days and later, when we sampled CE goldfish for transcriptomic analysis) of the CE goldfish, retina (including pigment epithelial cells and photoreceptors) was invaded and replaced by phagocytes and glial cells [5]. This is in line with our finding that melanogenesis, phototransduction, and phagosome pathways were affected by CE mutation.

Other than eye expansion and retina degeneration, we suggest that changes may also take place in the corneal of the CE goldfish. Compared with the transcriptomic analysis aiming for the TE goldfish [13], our study found that inflammatory reactions including Toll−like receptor signaling pathways were exclusively activated in the CE goldfish. Together with down regulations of keratinocyte and cornea related processes in the CE but not TE goldfish, a possible scenario is that, the corneal keratocytes, in the eyeballs of CE suffered heavy injuries which triggers inflammation and transformation of keratocytes into extracellular matrix (ECM) components secreting myofibroblasts [15]. At the same time, less keratocytes means less keratan sulfate and results in failure to maintain corneal hydration [16]. Terms or pathways related to the above processes were enriched in our transcriptomic analysis (cornea development in camera-type eye, regulation of water loss via skin, ECM-receptor interaction, et al.). Alternatively, PPAR signaling pathway could be the direct cause of inflammation and water loss [17] instead of keratocytes. The clinic characters of the cornea in the CE goldfish will be investigated in our future studies. What is more important, the direct cause of eyeball turning in the CE goldfish remains unknown.

Although *LRP2* is the only DEG with obvious functional mutations that matches the criteria of CE candidates, other DEGs should not be excluded since SNPs in the intergenic region could also be functional. Unfortunately, the lack of a functional motif or conservation data base for goldfish hinders the discovery of these functional mutations. As shown in **Fig 6B**, like *LRP2*, other DEGs (*FRZB*, *NEUROD1*, and *KRT18*) might also play a central role during eye development. *FRZB* (frizzled related protein, also known as *SFRP3*) is a Wnt signaling inhibitor [18] and evolutionally conserved in the vertebrates [19]. In a study of human ophthalmic disease Age-related Macular Degeneration (AMD), *FRZB* was identified as a mechanistic player in geographic atrophy which is a form of AMD and characterized by patchy degeneration of the retinal pigment epithelium and the photoreceptors [20]. *NEUROD1* (also known as *NEUROD*) is a basic helix-loop-helix transcription factor critical for regulating neuronal cell cycle, *NEUROD* knockdown in zebrafish prevented photoreceptor maturation and regeneration [21]. Furthermore, *NEUROD* in zebrafish was expressed rhythmically in differentiating photoreceptors and also in adult retina [22]. *KRT18* encodes a type I keratin which is expressed in wide range of tissues in humans [23]. Recently, knockdown of *krt18a.1* (a duplicated *KRT18* gene) in zebrafish suggests that it contributes to early development of ocular neural crest cells and corneal regeneration in adults [24]. Although *CERKL* and *klhl41a* were not predicted to interact with other DEGs, other studies provide clues for them to be possibly functional during the development of CE. Knockdown or knockout of *CERKL* (ceramide kinase-like) in zebrafish were generated, and degeneration of photoreceptor and apoptosis of retinal cells were repeatedly reported [25–27]. While small eyes were observed in only one of *CERKL* knockdown zebrafish [26]. *klhl41a* (kelch-like family member 41a) was highly expressed in eyes at 1 dpf but not 2 dpf of zebrafish embryo, and knockdown zebrafish also exhibit smaller eyes along with leaner bodies and pericardial edema [28]. *ITPRID2* and *AGR3*, were predicted to interact with *FRZB* and *KRT18*, respectively (**Fig 6B**), but the relationship between *ITPRID2* and eye development was not reported yet. Expression of *AGR3* was found by single-cell transcriptomics in a cluster of cells that were presumably proliferating corneal epithelial cells [29]. In addition, over-expression of *AGR2* generated enlarged eyes in *Xenopus* embryo [30], but whether *AGR3* has the same effect is unknown. *LOC113071285* is an uncharacterized protein and thus no more information was found.

Taking together, among the above mentioned DEGs, *FRZB*, *NEUROD1*, and *KRT18* were only involved in retina or cornea related processes, while *klhl41a* knockdown induced zebrafish with smaller eyes but no retina degeneration was reported. Therefore, these genes are less likely to be the cause of CE. Although retina development and eye size were modified by knockdown or knockout of *CERKL* in zebrafish, small eyes were not constantly observed. In addition, *CERKL* was upregulated in CE goldfish (**Fig 6A**), suggesting that its causality to CE needs to be further investigated. In contrast, if the same as *AGR2*, *AGR3* over expression can enlarge eyes, it could not be responsible for CE in goldfish since it was down regulated in CE eyeballs (**Fig 6A**).

Lastly, *LRP2* remains to be the top candidate, not only because of its functional mutations and differential expression, but also for its reported functions. Multiple *LRP2* knockout mice lines showed enlarged eyes and less retinal cells, but normal intraocular pressure [31–33]. Mutations in human *LRP2* lead to Donnai-Barrow and Facio-oculo-acoustico-renal (DB/FOAR) syndrome, characterized by buphthalmia (protuberant eyes), high-grade myopia, *et al* [34,35]. Premature stop codon of *LRP2* in zebrafish induced naturally or artificially, similar to the causal mutations identified for the TE goldfish and CE goldfish in this study, also exhibited enlarged eyes, retina degeneration, elevated intraocular pressure, and severe myopia [36–38]. Furthermore, signs of phagocytes in the retina of *LRP2* deficient mice were detected [39], resembling those in the CE goldfish [5].

In conclusion, the gene mapping in this study suggests that coding mutations of *LRP2* gene is responsible for CE in goldfish, which gains our knowledge regarding the genetics and evolution of TE and CE at the molecular level. Moreover, transcriptomic analysis provides clues regarding to the cellular processes contributing to the CE phenotypes.

## Materials and Methods

### Ethics statement

All experiments were conducted according to Guidelines for Experimental Animals established by the Ministry of Science and Technology (Beijing, China). Animal experiments were approved by The Science Ethics Review Committee of Beijing Academy of Agriculture and Forestry Sciences (Beijing, China) (approval number: Baafs20240901).

### Animals and tissue collections

In a CE × CE mating, 3 NE offspring out of 1,277 offspring (others were CE or TE phenotypes) were chosen to build the mapping population (NEM, NEF1, and NEF2). Then the two crosses, NEM × NEF1 and NEM × NEF2 were executed. Cross1 (NEM × NEF1) produced 309 goldfish, while 634 goldfish were obtained from Cross2 (NEM × NEF2). In Cross1, 59 goldfish with standard CE phenotype (the eyeballs were 90° turned upwards) and 80 goldfish with NE phenotype were collected for their tail fins; in Cross2, 64 standard CE goldfish and 80 NE goldfish were also collected for their tail fins. The tail fins for the 3 parental goldfish, NEM, NEF1, and NEF2 were also collected. Those goldfish were kept in a glass aquarium measuring 1.2 meters in length, 0.6 meters in width, and 0.45 meters in height. The water was static and 35 cm in depth, using groundwater that has been aerated for more than 48 hours. The aquarium was indoor but experiencing the natural temperatures and lighting through the year. During the experiments, the amount of feeding was adjusted based on the body conditions of the fish to keep them healthy. DR900 Multiparameter Portable Colorimeter (Hach, USA) was used to monitor water conditions, ensuring that the pH values were between 7.0 and 8.4, dissolved oxygen was from 9.70 to 7.70 mg/L, nitrite was less than 0.02 mg/L, and ammonia nitrogen was less than 0.15 mg/L. Water was changed frequently depending on the quality conditions. Phenotypes of the eyes of these goldfish were recorded after 12 months of age.

Eyeball tissues (without eyelids or surrounding connective tissues) for RNA-seq were collected for 3 NE goldfish from a fish farm at Jiangsu Province, China, and 3 CE goldfish from a fish farm at Beijing, China. These 6 goldfish were purchased at 6 months of age, then kept in the same condition as described above for 8 months before euthanizing and sampling. Tail fins of these 6 goldfish were also collected for DNA extraction.

In addition, tail fins of goldfish were collected from 5 independent fish farms located in different regions of China, i.e., Anhui (Ah), Beijing (Bj), Hebei (Hb), Jiangsu (Js), and Shanghai (Sh) (n=10 for each farm), and 10 NE from Jiangsu Province for diagnostic tests.

### DNA extraction and sequencing

The above mentioned 356 fin samples were extracted for DNA via CTAB with modifications [40]. Invitrogen Qubit 4.0 (Thermo) and 0.8% agarose gel were applied for the quality control of DNA. Four DNA pools were constructed according to the shared phenotypes or crosses (Pool_CE1, Pool_NE1, Pool_CE2, and Pool_NE2). Each sample contributed 15 ng of DNA to the pool. For WGS, the 4 DNA pools and 3 parental goldfish DNA samples were prepared by TruSeq DNA PCR-free prep kit (Illumina, USA) which built the libraries with the insertion size of ∼450 bp, and followed by the quality control accomplished by 2100 Bioanalyzer (Agilent Technologies, USA) via High Sensitivity DNA Kit (Agilent Technologies). After quantification of the libraries by QuantiFluor (Promega, USA) via Quant-iT PicoGreen dsDNA Assay Kit (Thermo Fisher Scientific, USA), the qualified libraries were 2×150bp paired-end sequenced by Illumina NovaSeq with 30X coverage (the 4 offspring pools) or 10X coverage (the 3 parental samples). The 50 CE goldfish samples were also prepared and sequenced individually with 4X coverage. All the sequencing was carried out by Personalbio Technology Co.,Ltd (Shanghai, China) and raw data was deposited in the Genome Sequence Archive [41] in National Genomics Data Center (NGDC) [42], China National Center for Bioinformation / Beijing Institute of Genomics, Chinese Academy of Sciences.

### Alignment of WGS data and calling of SNPs, InDels, and SVs

The raw sequencing data were analyzed and quality controlled by FastQC (version 0.12.1). Then the clean data were aligned to the goldfish reference genome [11]. All FASTQ clean data were aligned to the reference genome using BWA-MEM (version: 0.7.12-r1039) [43] with default parameters. Then the SAM files were sorted and converted to BAM files by SAMtools (version: 1.17) [44].

After the alignments, SNPs and InDels were called with GATK HaplotypeCaller 3.8 but no filtration was applied, in case of excluding possible causal mutations, and they were annotated by ANNOVAR (version 2019-10-24) [45]. Structural variants were called with Lumpy (version: 0.2.13) [46].

### Diagnostic test for the reported TE causal mutations

PCR protocols with a 3-primer system were designed to genotype the ∼13 kb insertion in the 45^th^ intron of *lrp2aL* gene [9]. The primers are LRP2_Exon45_F (5’-GCAGTGATGGTTCGGATGAG-3’), LRP2_Exon46_R (5’-AACTGGTCGGAGTTGCAGGT-3’), and gFV-1_R (5’-CCCAGTGAGACACGATTGGA-3’, or gFV-1_F: 5’-AGATTGCCTTTGCTGGTTTGA-3’). LRP2_Exon45_F and LRP2_Exon46_R can amplify a 1,710 bp fragment when the 13 kb insertion was absent, while LRP2_Exon45_F and gFV-1_R (or LRP2_Exon46_R and gFV-1_F) can only amplify fragments when the allele with the ∼13 kb insertion exists (since the exact insertion site is unknown, the precise PCR product sizes are not clear but should be less than 1,710 bp). In addition, a 2-primer system PCR was also carried out using primers LRP2_Exon45_F and LRP2_Exon46_R, for Sanger sequencing of the intron 45 of *LRP2*. Samples used for sequencing the 45^th^ intron of *LRP2* gene were 2 randomly selected CE goldfish in the offspring of Cross1 and another 2 random CE fish in the offspring of Cross2.

The 3-primer PCR systems include 10 μl of 2×Tap Plus PCR MasterMix (Solarbio, Beijing, China), 0.5 μl of each primer (10 μM concentration), 1 μl of genomic DNA (50 ng/μl concentration), and ddH_2_O added up to 20 μl. The protocol was used for all the PCR of 94°C for 3 min, 30 cycles of 94°C for 30 s, 60°C for 30 s, and 72°C for 1 min each, followed by 72°C for 10 mins. Amplifications were carried out on a Veriti 96 well thermal cycler (Applied Biosystems, Thermo). PCR products were assessed by agarose gel electrophoresis or Sanger sequenced (Tsingke Biotechnology, Beijing, China).

### Identification of candidate regions and mutations

The BAM files for Pool_CE1 and Pool_NE1 were used to calculate pair-wise F_ST_ values with Popoolation2 in a sliding window approach with window size of 50 kb and step size of 10 kb. Allele frequency differences (dAF) between the 2 pools were also calculated by Popoolation2. Pool_CE2 and Pool_NE2 were analyzed according to the same procedure. The F_ST_ values were Z-transformed and genomic regions with ZF_ST_ values higher than 11 were defined as candidate regions (the sum of high ZF_ST_ regions detected in Cross1 and Cross2). Within the candidate regions, variants with dAF larger than 0.5 while being heterozygous in both the parents were considered as candidate mutations. Next, all candidate mutations that were fixed in the 50 individual WGS data from 5 different fish farms were screened out.

For each of the candidate regions, the pair-wise genetic distances between the 52 CE goldfish libraries (2 pools and 50 individuals) were evaluated by the following steps: 1. The QC of raw output of GATK HaplotypeCaller was consist of “QD < 2.0 || FS > 60.0 || MQ < 40.0 || MQRankSum < -12.5 || ReadPosRankSum < -8.0” of filter-expression option in GATK VariantFiltration for SNPs, while indels were excluded; 2. The filtered SNPs were phased by beagle (version: 5.1); 3. Python script distMat.py (https://github.com/simonhmartin/genomics_general) was applied to calculate the pair-wise genetic distances between all the 52 libraries. Any of the 50 individual samples showing genetic distance with Pool_CE (1 or 2) larger than 0.1, were excluded for the screening of target mutations for this region, since these sequences were considered originated differently from our mapping population and thus not sharing the same causal mutation.

All the SVs within or flanking the target region (1 Mb upstream and 1 Mb downstream), plus candidate coding SNPs and Indels, were selected for further analysis. All the selected SVs were double-checked for their reliability and allelic frequencies in the 7 libraries from the mapping populations by viewing the BAM file to sift out the ones that were reliable and also matches the criteria “heterogeneous in the 3 parental samples and 2 NE pools but homozygous in the 2 CE pools”.

### RNA extraction and transcriptomic analysis

For RNA-seq analysis, total RNA was extracted from the 3 NE and 3 CE goldfish using Trizol (Invitrogen, Carlsbad, CA, USA) according to the manufacturer’s protocol. Library products were prepared and NovaSeq 6000 platform (Illumina) was applied by Personalbio Technology Co.,Ltd (Shanghai, China). Cutadapt (version: 1.15) [47], Hisat2 (version: 2.0.5) [48], and HTseq (version: 0.9.1) software were used for quality control, aligning to the goldfish reference genome [11], and counting for read number for each gene. Next, DESeq (version: 1.30.0) was applied for detecting differentially expressed genes between NE and CE groups. The criteria for DEGs are |log2foldchange| > 1 and adjusted *P*-value < 0.05. Then GO and KEGG enrichment analyses for the DEGs were done by the clusterProfiler package (version: 3.4.4) of R. The same protocols were applied to all the raw short read RNA-seq data uploaded by Du et al [12] (PRJNA558211).

Interaction network and gene prioritization for candidate genes were predicted by the online platform GeneMANIA (https://genemania.org/) [49].

### Diagnostic test for the candidate mutation for CE

To genotype the premature stop codon of *LRP2* (chr9: 28,575,379), primers were designed: LRP2_Exon38_F (5’-GGACCACCGAGACGGATACA-3’) and LRP2_Exon38_R (5’-TGGGATGGCGAAGCAGA-3’). The amplicon is 361 bp and the same PCR protocol was applied as mentioned above. Both forward and reverse primers were used for Sanger sequencing for double checking. The Sanger sequenced goldfish were the CE offspring in our mapping population (n=11), which the eyeballs were 15° turned upwards (n=1), 30° turned (n=2), 45° turned (n=1), 60° turned (n=2), 70° turned (n=1), one eye 80° and another eye 60° turned (n=1), one eye 90° and another eye 60° turned (n=1), one eye 90° and another eye 70° turned (n=2). Besides, the offspring with slightly protuberant eyes (no turning of eyeballs, n=5), or NE (n=6) were also selected for Sanger sequencing.

## Acknowledgments

The authors acknowledge Dr. Zhen Huang from College of Life Sciences, Fujian Normal University, Fuzhou, China who provided the goldfish reference genome for this study.

## Supporting information

**S1 Table. Candidate regions of CE in goldfish through the comparisons of offspring in mapping populations by pooled WGS.** Candidate regions were defined as ZFST larger than 11. Firstly, candidate regions in Cross1 and Cross2 were defined separately (middle and right panel in the table, respectively), then they were combined into the candidate regions in the left panel in the table.

**S2 Table. The identity regions shared by the 50 CE goldfish from 5 different populations.** The data is coordinated with **Fig 4**, showing that no region was shared by all the samples.

**S1 Fig. The SVs detected in the intron 45 of *LRP2* gene.** “TE?” in this figure indicating that these SVs (SV1 and SV3) could be transposon elements because of the surrounding reads (mates mapped in a different chromosome, and secondary mapping). The allelic frequencies of these 5 SVs matched the expected pattern for causal mutations of CE except SV2 which is almost 100% in 4 DNA pools. Sanger sequencing has confirmed that SV1 is a 33 bp sequence (chr9:28,583,267-28,583,299) replaced by a 116 bp sequence, and also the previously reported ∼13 kb insertion is absent.

**S2 Fig. Enrichment analysis of the differentially expressed genes (DEGs) between eyeballs of NE and CE goldfish via RNA-seq.** Top 20 GO terms (*A*) and top 20 KEGG pathways (*B*) significantly enriched for all the DEGs. In each of *A* or *B*, the *P*-value increases as the terms or pathways are presented from the bottom to the top.

